# Imaging microvasculature network evolution and neurodegeneration with precise photothrombosis approach

**DOI:** 10.1101/2021.11.29.470313

**Authors:** Liang Zhu, Mengqi Wang, Yin Liu, Weijie Zhang, Hequn Zhang, Anna Wang Roe, Wang Xi

## Abstract

In the cerebral cortex, the vasculature plays important homeostatic functions, especially at the highly connected complex capillary networks. The association of focal capillary ischemia with the neurodegenerative disease as well as the laminar vascular dynamics have prompted studies of vascular micro-occlusion via photothrombosis. However, technical challenges of this approach remain, including increased temporal precision of occlusion, increasing the depth of vascular occlusion, understanding how such micro-occlusion impacts local blood flow, and ultimately the neuronal effects of such changes. Here, we have developed a novel approach that employs ultra-fast multiphoton light to induce focal Rose Bengal-induced photothrombosis. We demonstrated induction of highly precise and fast occlusion of microvessels at various types and depths. The change of the microvascular architecture and hemodynamics after occlusion revealed the autoregulation and significant difference between upstream vs downstream in layer 2/3. Further, we found that micro-occlusion at two different layers within the same vascular arbor results in distinct effects on the acute flow redistribution mechanism. To examine neuronal effects of such micro-occlusion, we produced infarct of capillaries surrounding a labeled target neuron and found this induces dramatic and rapid lamina-specific degeneration in neuronal dendritic architecture. In sum, our technique enhanced the precision and power of the photothrombotic study of microvascular function. The current results pointed to the importance of laminar scale regulation within the microvascular network, a finding which may be relevant for models of neurovascular disease.

## Introduction

The brain vascular system dynamically supplies region-specific oxygen and glucose while also transporting their metabolic waste products for eventual elimination [1]. The microvascular network within the brain parenchyma is uniquely positioned to detect the neuronal and astroglia signals to modulate the blood flow [2]. The challenge to investigate dynamic microvasculature change is due to the complex flow dynamic and network connectivity *in vivo*. How the capillary network maintain homeostasis is important for the survival of neighboring neuron and astroglia. In the cerebral cortex, the highly integrated microvascular network functionally correlates with laminar neural activity. The type of neuron computation connection performed across cortical layers strongly influences the organization of the cortical microvascular topology, indicating the oxygen tension difference in cortical layers [2, 3]. There has been a long-standing interest in how the brain regulates its blood supply, driven not only by the desire to gain a better understanding of the adverse effects of cerebrovascular insufficiency but provide an early indication of regional activity changes in cerebral blood flow (CBF) may provide a window on brain function. Therefore, the layer-specific vascular micro ischemia model is needed to understand the precise microvascular regulation mechanism of the blood supply across laminas.

Small cerebral infarcts, i.e. microinfarcts, are common in the aging brain and linked to vascular cognitive impairment. However, little is known about the acute growth of these minute lesions or their effect on the blood flow in surrounding tissues. These microvascular dysfunctions have contributed to neurodegenerative disorders such as vascular dementia, Alzheimer’s disease, ‘silent’ stroke, ALS, and psychiatric manifestations such as autism spectrum disorder [4–8]. How the micro ischemia interacts with these neurodegenerative diseases may clarify its clinical relevance. Many efforts have been taken to develop the selective target cerebral microvascular lesion, which resides below the surface of the brain. Produce focal ischemia via occlusion mediated with a suture or ligation [9–11], thrombotic blood clot emboli [12–14], dye-induced photothrombosis (e.g., using Rose-Bengal or erythrosine B) [15–18], and occlusion mediated through endothelin-1 [19–21]. Furthermore, thrombosis can be introduced by inducing focal occlusion in a single microvessel with an infrared laser [22] and reverse cerebral ischemia with magnetic nanoparticles [23]. The small vessel micro-occlusion also can be produced brain-wide through fluorescent microspheres [24]. Hence, it remains a challenge to induce focal micro-ischemia models with precise targets with capillary levels in the brain to control the ischemia sizes and depth.

To understanding of the relationship between cortical structure, function, and blood flow in the capillary level in health and disease, here, we combine the dye-induced photothrombosis with ultra-fast multiphoton light to induce the precise occlusion of the various size of vessels deep to 800μm depth in layer 6. In combination with the two-photon microscopy, we can image the dynamic microvascular network autoregulation during the smallest capillary ischemia in minutes. This precise focal ischemia was also used to image the laminar-specific infarct in the Thy1-GFP mice to reveal the single neuron degeneration changes. Using this precise target micro occlusion method, we ask questions about the basic capillary function in the microvascular network resistance and its relationship to neurodegeneration. 1. Does the laminar difference topology connections result in different microvascular autoregulation mechanisms? 2. How does the microvascular network evolution after the single capillary occlusion? 3. Does the capillary occlusion cause regional neurodegeneration?

## Result

### Establishing a precise single vessel photothrombosis *in vivo*

To create the focal ischemic, we used a 1070nm femtosecond laser to photoactivate Rose Bengal in cerebral vasculature within 5 minutes, due to Rose Bengal metabolism (Fig. 1a.c). The stimulation laser used a galvo-galvo system which was configured with a customized imaging system (Fig. 1a). Firstly, we used a fluorescent board to prove the reliability of the laser stimulation pattern. After a single stimulation paradigm was delivered, a cleanly burnt spiral path appeared in the chosen ROI (Supplementary Fig.1). Then, we tested the efficiency of the nonlinear optical excitation of Rose Bengal on the somatosensory cortical vasculature in head-fixed mice with TPLSM under isoflurane (1-3%) (Fig. 1 a,b). The vessels were labeled with fluorescein-dextran (2 MDa, FITC or TRITC) to verify structural and functional changes after photothrombosis. We mapped a three-dimensional vascular architecture in the vicinity of the targeted vessel during baseline using a stack of high magnification images (Fig. 1d). The 1070nm femtosecond laser was modulated to a point-by-point, spirally patterned stimulation sequence that photoactivated Rose Bengal in a tight focal point of a single targeted vessel, which repeated stimulation caused aggregation of blood platelets and eventual stagnation of blood flow (Fig.1f). Our example showed a pial trifurcating arteriole, whose uniform baseline flow evidenced by streaks in the line-scanned image (Fig. 1e). The clot formation was imaged in real-time, demonstrating the flow alteration of the arteriole (Figure 1d-f; Supplementary Movie 1). Measurements of the changes of speed and direction of RBC flow after the occlusion formation in the target and neighboring vessels indicated that the occlusion effectively caused redistribution in vessels that lie immediately downstream (Fig. 1d, e). Conversely, we proved no occlusions and collateral damage were induced in adjacent vessels near the targeted irradiated area with identical flow post-stimulation. Further, laser stimulation of the same paradigm without Rose Bengal showed no sign of thrombosis in the targeted vessel (Supplementary Fig. 3).

**Figure 1.**
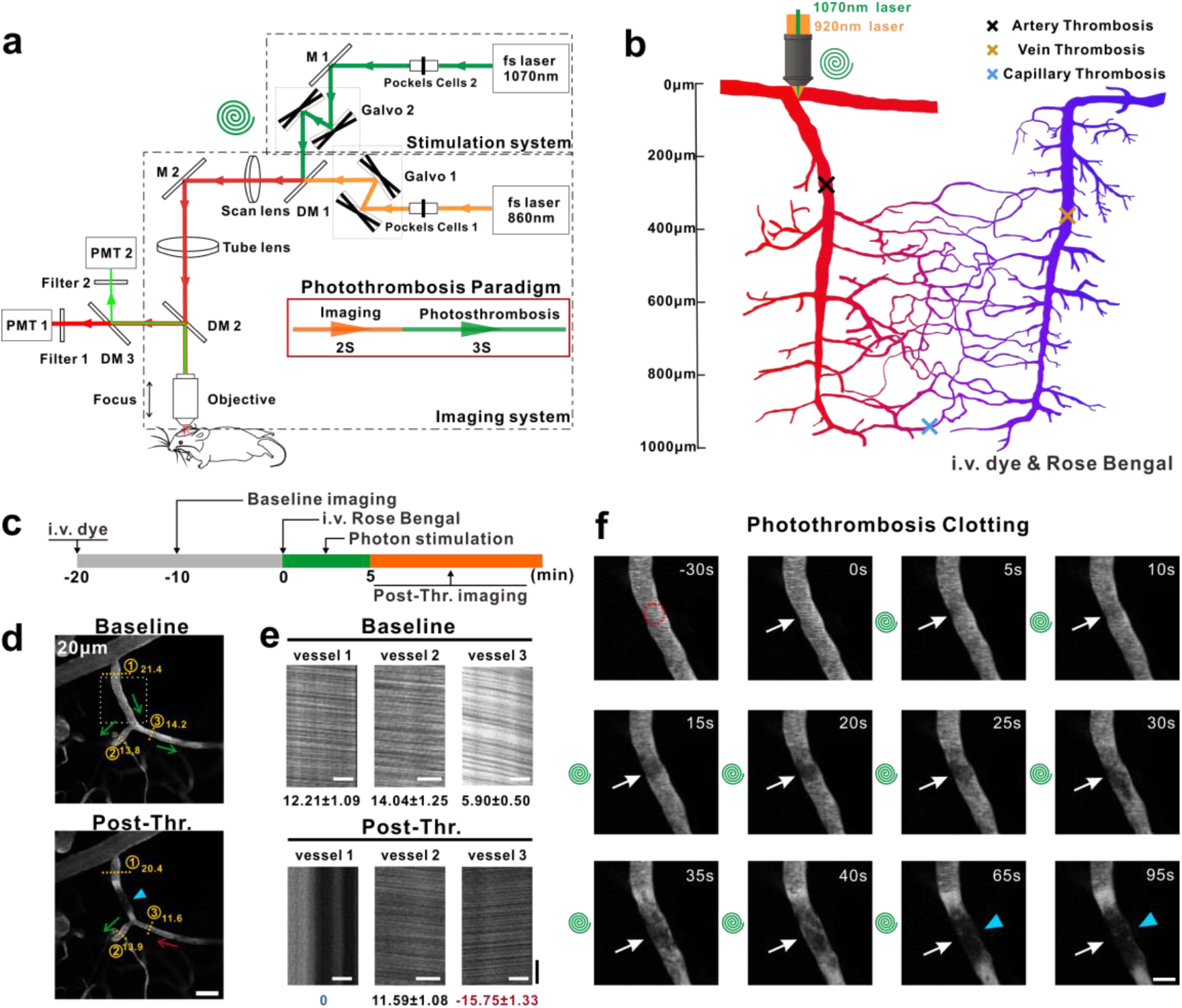
Simultaneous single vessel nonlinear photothrombosis with two photon imaging paradigms. a. Schematic of the nonlinear photostimulation configuration with the two-photon laser scanning microscope system. The optical paths for two-photon imaging (920nm, orange) and targeted photothrombosis (1070nm, green) were sketched. Two beams of light enter the microscope system through two sets of galvanometers separately. The red rectangle indicates the stimulus paradigm. b. Illustration of nonlinear laser induced photothrombosis. Schematic depicting zones in cortical microvasculature. 1070nm laser light was focused through an objective and directed into the target vessel of different depths and types (marked as X), that photo-activated the intravascular Rose Bengal dye and initiated a clotting cascade. c. Experimental time course for photothrombosis of the targeted vasculature imaging. d. An example of a single vessel photothrombosis in the cortical surface. (1, 2, 3 indicated the different branches of the artery; dotted crosssection lines showed the diameter of each segment. Blue arrowhead referred to the irradiated site, green arrows showed the baseline blood flow direction, and red arrow indicated reversed flow, notably in post thrombosed vessel 3). Notably, the direction of blood flow in the downstream in fig1c vessel 2 changed. Scale bar, 50μm. e. Line-scan measurement of the blood flow speed of the arteries in figd before and post occlusions. Negative values in red indicate the vessel with a reversed flow after clot formation. Unit, mm/s; horizontal scale bar, 20μm; vertical scale bar, 0.01s. f. Snapshots of real-time planar imaging during the intravascular clot formation, enlarged from the white rectangle in fig1c. The red dotted circle in the first snapshot showed the area to be affected by the focal photothrombotic irradiation for a targeted vessel. The green spiral in between snapshots indicated the path of one pass of laser stimulation. The white arrows indicated the eventual formation of a thrombus clot, with a stable thrombus (blue arrowheads) formed at the 65th second. Scale bar, 20μm.

### Demonstration of single deep capillary and simultaneous multi-capillary occlusion

Previous studies have established some methods to produce targeted single vessel disruptions in the brain cortex [22, 25]. However, technical challenges remain on unwanted stimulation to nearby microvessels due to a reduced spatiotemporal accuracy with an increased depth. Our refined method provided restricted parameters to more accurately and quickly occlude arteries, veins, and capillaries of different sizes and depths with less/no collateral damage (Fig.2, Supplementary Fig. 4). The pial artery usually requires more substantial laser power and much more stimulation time due to the high flow speed and excellent elasticity of the vessels. Notably, the pial vein, which has much slower blood flow and worse contractility, was easier to form a clot using fewer stimulation sequences with less energy (Table1). In addition, we observed the flow stagnated in the targeted vessel last for 2 hours and the flow redistribution and dilation/constriction in upstream and downstream vessels (Supplementary Fig. 4, Supplementary Movie 2).

We demonstrated the single and multi-capillary photothrombosis models at different depths without collateral damage (Fig. 2). For single capillary, we successfully induced occlusion at 150μm, 300μm, 500μm, and 770μm (Fig. 2a-h). The upstream of the occluded vessel showed slight vasodilation compared to baseline, and the reverse holds true for downstream (Fig. 2a, c, e, and g). For multi-capillary occlusion, three individual capillaries at 200μm below the pial surface were stimulated with a 3-second interval each followed with 2 seconds of imaging (one sequence). After several stimulus sequences, occlusion was observed in each capillary simultaneously and is supported by line-scan blood flow data (Figure 2i, j, Supplementary Movie 3). Further, we showed an accurate single capillary occlusion at an extreme depth (815μm) without collateral damage (Figure 2k-p). We examined the metrics of three layers/segments above the target capillary via a 3D reconstructed image and showed normal flow (Supplementary Fig. 5) and architecture before and after the thrombosis (Fig. 2k-n). A clot was observed in the targeted capillary, while the flow velocity decreased by 65% with a slight dilation in upstream vessel, and a complete stasis in the downstream vessel (Fig. 2o, p).

**Figure 2.**
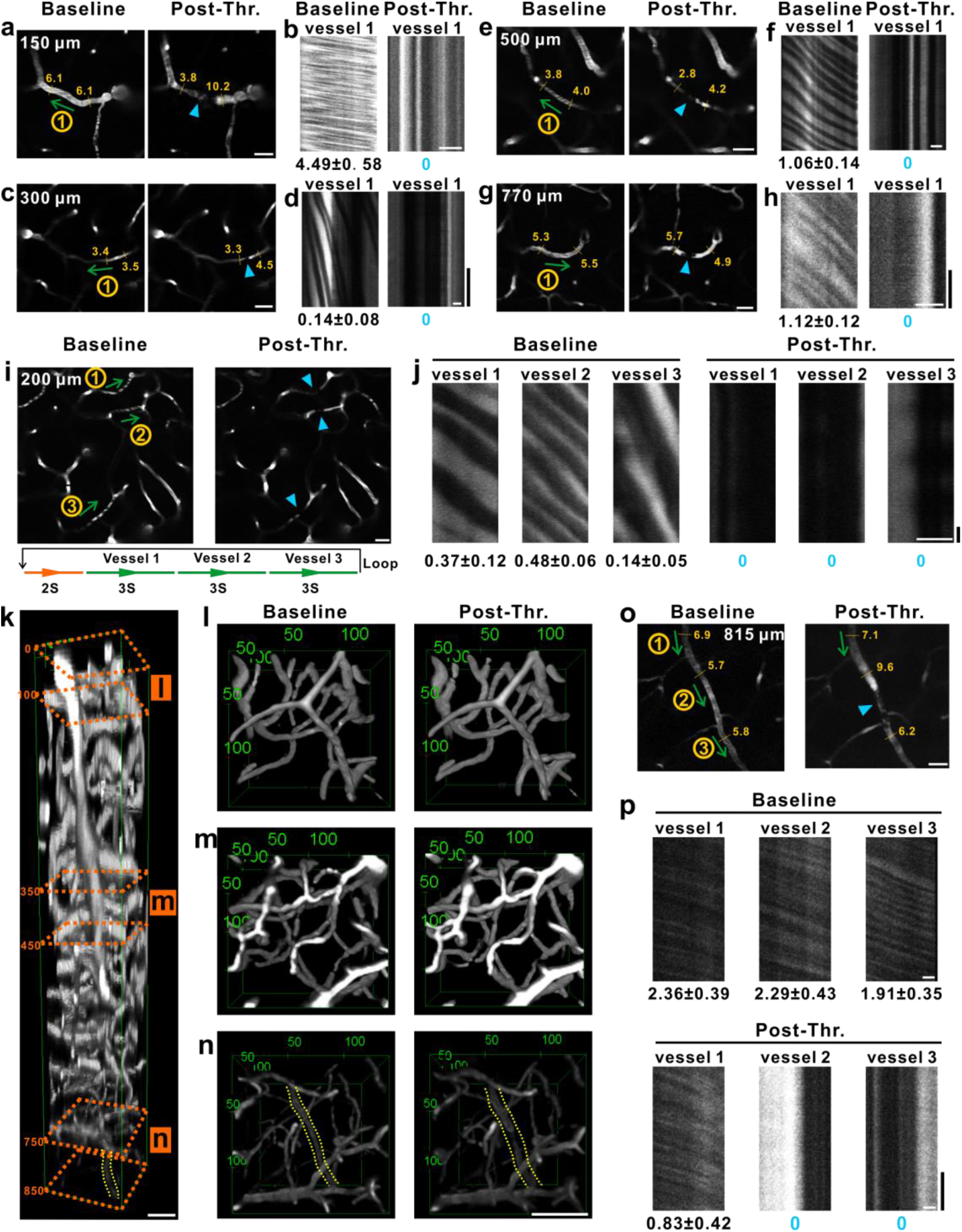
Demonstration of fast and precise photothrombosis of single or multiple capillaries at different depths. Capillary imaging (A, C, E, G, I, O) and respective line-scan (B, D, F, H, J, P) result on pre and post photothrombosis at corresponding depths (150 μm, 300 μm, 500 μm, 700 μm, 200 μm, 815 μm). Circled Ⓝ ^th^ number indicated targeted capillary or capillary segment. Yellow number with corresponding dashed line indicated capillary diameter up or downstream (Flow direction: green arrow) the site of pre (Left) or post occlusion (Right, blue arrowhead). Scale bar, 20μm. The number below each line-scan indicated flow velocity of targeted Ⓝ^th^ vessel pre and post thrombosis. Unit, mm/s. horizontal scale bar, 20μm; vertical scale bar, 0.01s. Additionally, the diagram below ‘I’ indicated the temporal paradigm of light-induced thrombus stimulation. Perspective view (K) of a reconstructed in vivo image stack (0-850 μm). Segmented bird’s eye view (L-N) of their respective depth range (Orange marquee, 0-100 μm, 350-450 μm, 750-850 μm). The yellow dotted line (detailed in O, P) indicated the target capillary. Scale bar, 50μm.

### Laminar-specific capillary network flow redistribution after pre-capillary arteriole occlusion

In the cortex, the distribution of neurons and consequently the metabolic needs vary over the depth of layers. Thus, the distinct layer-specific vasculature varies accordingly to support the demand, with the highest vascular density and topology connections found in different layers of the primary sensory cortex in mice [26–28]. To test the heterogeneity of laminar-specific capillary network flow, we induced photothrombosis of the pre-capillary arterioles in layer2/3 and layer 4 within the same vascular arbor, which were the original and most important source of the downstream capillary network supply within these laminas[4–6] (Fig. 3a, b). In this case, two pre-capillary arterioles in lamina 2/3 and laminae 4 were selected for photothrombosis and 3D reconstruct of the downstream vessel trees were measured. We divided and measured the 1-to-5th order capillaries extended from descending artery whose only source was the target pre-capillary arteriole (Fig. 3d). The flow velocity of each vessel segment and the architecture were measured by TPM imaging before and after occlusion. (Fig. 3c).

**Figure 3.**
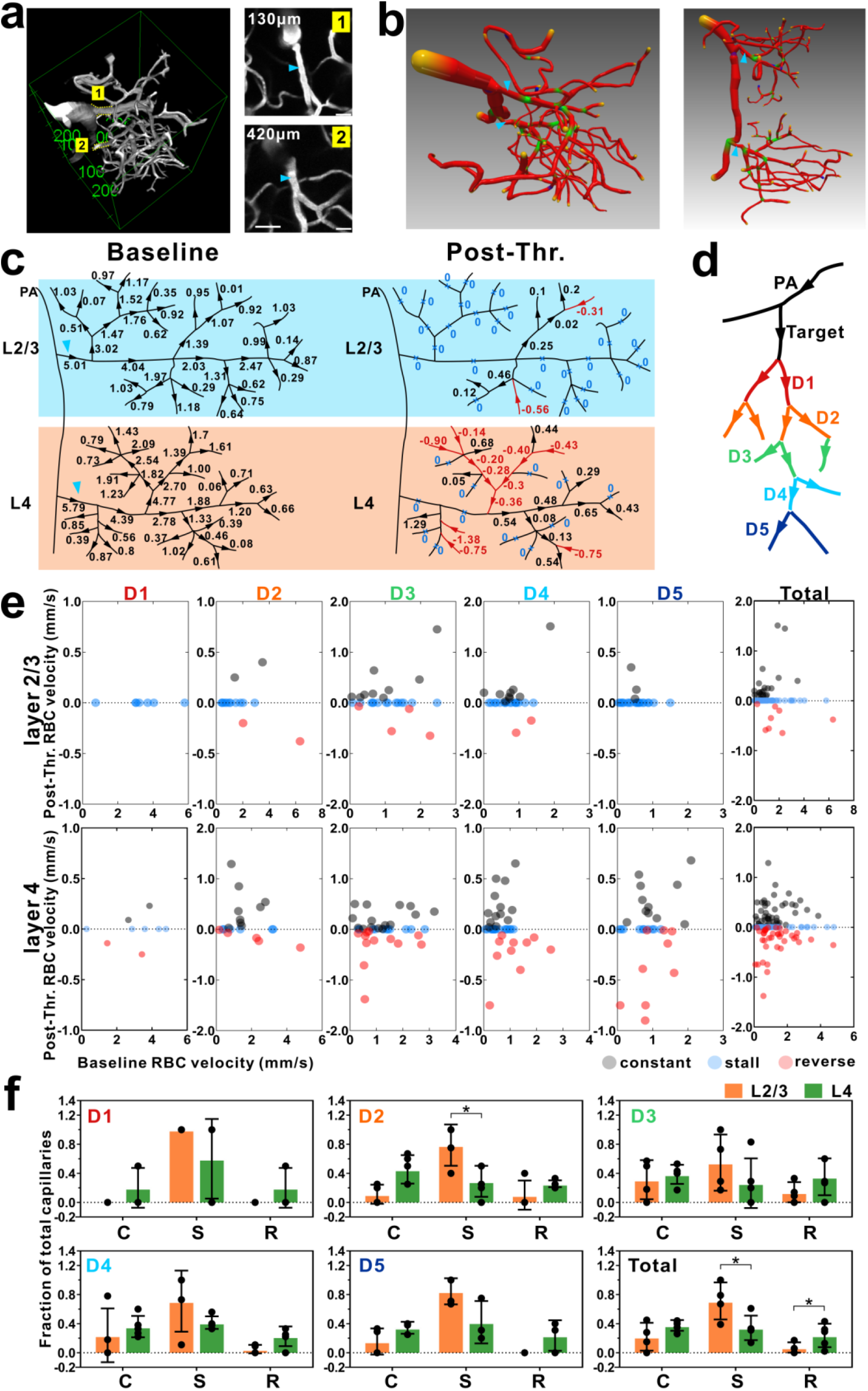
Heterogeneity of layer-specific capillary network vasodynamics response to acute pre-capillary arteriole photothrombosis. a. Left shows an example 3D TPM reconstruction of a single penetrating artery to capillaries network with their spatial position and sequential branch orders relative to the targeted pre-capillary arteriole. Right displays enlarged images obtained from the stack at their respective depth (130um and 420um) with the pre-capillary arteriole (8μm) occlusion (blue arrowhead). Scale bar, 20μm b. The rendered and reconstructed bird’s eye and side perspective views of the target vascular network of Fig 3a. Red indicates individual vessel segment, green shows the branch nodes, yellow refers to the capillary terminals, and the blue arrows correspond to the two pre-capillary arterioles. c. Flow change of the microvascular network induced by the occlusion of the two-arteriole indicated by the blue arrowhead. The blood flow velocity measurement of two vessel networks before and after the thrombus was depicted. (Black and red arrows indicate the direction of blood flow, and the blue X marked the stalled capillaries). d. Classification scheme of the vessel network. (The color coding indicates the branch orders, D1 to D5 refer to downstream vessels). e. Three categories of blood flow velocity changes of each branch order before and after the arteriole clot. Dots and coloring indicate the flow status of each capillary (Blue indicate stasis; black indicate a continuous flow; red indicate a reversed flow. N= 5 mice, T= 237 vessels). f. Column plot of three categories of capillaries in fraction (C= continuous, S= stasis, R= reversed) compared between L2/3 to L4 for each of the respective branching orders post occlusion, the same capillary comparison without accounting for branching but grouped per individual mouse was included as “Total”. N= 5 mice, T= 237 vessels. Each data dot represented the fraction of each category of capillaries in one mouse. (P < 0.05)

In a total of 5 mice, we categorized the downstream vascular hemodynamics into 3 types based on the forms of flow after occlusion, including (1) constant (blood keeping flow), (2) stall, (3) reverse (opposite in flow direction), in 126 individual capillary segments (Fig. 3e). A capillary was categorized as “constant” when maintaining a flow of the same direction, while no RBC flowed over 5 minutes was defined as “stall”. All the 126 capillary velocity were plotted in Fig. 3e. All downstream branches of all laminas showed a significant reduction in blood flow, with most capillaries being stagnant, and a smaller proportion displayed either a continuous or a reversed flow (Fig. 3e, f). When comparing layers 2/3 and layer 4, the redistribution of blood flow was significantly different (Fig. 3f). In general, D1 branches at layer 2/3 were all stalled, whereas a few branches had a slight positive or negative flow at layer 4 (Fig. 3e). The D2 to D5 branches had different reactions in different layers. Blood flow reversal usually happened in D2 to D4 branches of layer 2/3, while more flow reversal occurred in all levels from D1 to D5 branches of layer 4 (Fig. 3f). Reversal flow usually came from the further downstream branches, whose original source were the adjacent individual diving arteriole or ascending venule. When the capillaries were grouped per individual mouse, the proportion of stagnant downstream branches in layer 2/3 was significantly more than that in layer 4 (0.713±0.255 for layer 2/3; 0.342±0.169 for layer 4; p = 0.0317,), while the proportion of reversed downstream branches in layer 2/3 is significantly less than that in layer 4 (0.073±0.071 for layer 2/3; 0.238±0.162 for layer 4; p = 0.0397) (Fig. 3e, f). The result indicated that in layer 4, the capillary network had a much better flow redistribution and collateral mechanism that the adjacent capillary network would immediately form a new flow pathway to support part of ischemic capillaries when blood flow was severely limited in the upstream branches due to the primary contributor. Conversely, due to a lack of collateral flow/contribution, the capillary network of layer 2/3 performed much worse. In addition, blood flow in layer 4 seemed to have robust homeostasis due to its vascular topology, which made it possible to produce a new acute redistribution mechanism when the flow source was insufficient.

### Capillary network flow autoregulation after single capillary occlusion

The capillary bed has plentiful topological connections to form a complex vascular network to perfuse the cortex in a local area. As reported, there were averaged 8 segments leading from diving arterioles and ascending venules [26]. Previous studies have also proved that clots targeted to single capillary caused highly reduce flow in the proximal 1-to-3rd order downstream vessel [22, 29], and clots targeted to single vessels in either surface or subsurface networks cause negligible ischemia or damage to tissues, which may due to a consequence of redundant flow pathways[22, 30]. However, the specific architecture and hemodynamic changes of upstream and downstream branches in this limited capillary network are still unknown. We used TPLSM to repeatedly image the patency and structure of subsurface capillaries within the receding peri-infarct tissues following occlusion of targeted capillaries in layer 2/3 (Fig. 4). And we investigated how changes in flow velocity and diameter of connected upstream and downstream vessels depend on the topological connection between individual segments and the occluded capillary.

**Figure 4.**
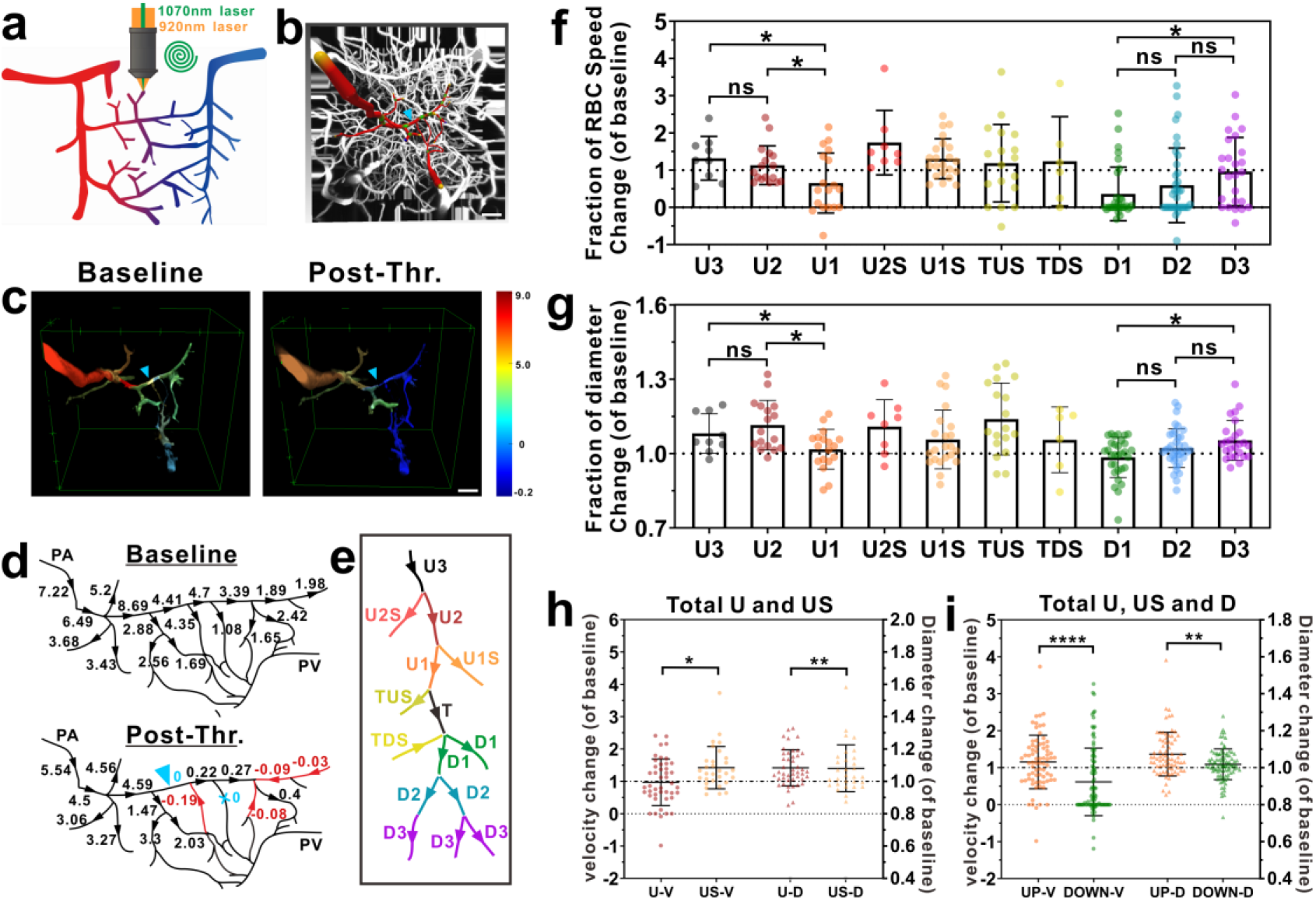
Capillaries network hemodynamic evolution after focal capillary occlusion. a. Schematic of the nonlinear photo-occlusion to the single capillary segment in the cerebral vascular tree. b. Example of a penetrating artery-to-6th branch order capillary branch connection. The 3-D rendering showed the traced artery-capillary orders with red color in the in vivo vasculature image stack side view (50-461μm). (Green: branch node; red: vessel segment; yellow and purple: segment ends) Scale bar, 50μm. c. Composite extracted tracing capillaries branch orders image corresponding to fig4b, e. The color corresponding to the color bar represented the blood flow velocity of different vessel segments. Blue arrowhead refers to the targeted clot. Color bar unit, mm/s. Scale bar, 50μm. d. Examples of flow changes that result from focal occlusion of a single parenchymal capillary. Baseline and post-occlusion blood flow speed measurements of the vascular network traced in panel b (blue arrowhead-marked corresponds to the capillary clot site. Color coding indicates the blood flow direction). e. Classification scheme of the vessel network order near occlusion site. Color coding indicates the branch order, U3, U2, U1, upstream branches; U1S, U2S, TUS, TDS, side branches; T, targeted capillary; D1, D2, D3, downstream branches. f,g. plot of the RBC speed change and diameter change of all orders of capillaries after occlusion. h. The RBC velocity and diameter change of all upstream branch and side branch segments before and after occlusion. i. The RBC velocity and diameter change of all upstream and side branch vs downstream branch segments before and after occlusion. (Error bars in all graphs are SD. Significant differences between groups *p < 0.05, **p < 0.01, ***p < 0.001, ****p < 0.0001.)

Firstly, we defined brain capillary network based on the flow direction and the topological branching order from the diving artery [31–33]. Each capillary segment was defined as a length of capillary extending between two branch points and divided into several orders (Fig. 4e). Capillaries selected for irradiation was defined as the target vessel (T). The vessels directly connected to target vessel were defined as upstream and downstream vessels (U1, D1), divided by direction of blood flow. Upstream vessels directly provided flow source to the target vessel and were grouped as one to three branches (U1-U3). In addition, parallel vessels that were directly connected to the target vessel or the upstream branches and shared the same source are called side branch vessels (U1S, U2S, TUS, TDS), which also divided by flow direction. Downstream vessels drained from the target vessel and were grouped as those immediately downstream (D1) and those downstream (D2 to D5) from the target. Forth to sixth branches were selected as the target vessels in our study, which ensured the measurement in similar vessel network. To quantify the targeted specific capillary network, the connection upstream and downstream network of the target capillary were imaged and traced, then extracted from the original vascular stack image, and assigned different colors. (A rotating movie of this image stack is shown in the Supplementary Movie 4, Fig.4c). We illustrated a typical example of flow changes (Fig. 4 c, d), in which flow was slowed in the upstream vessels, while flow was slightly reduced in a parallel vessel that shared the same source as the targeted vessel. In addition, several downstream branches reversed flow, accompanied by a decrease in the speed of the RBCs. Occlusion of single capillary generated a restrict range of ischemia volume in mouse cortex (Fig. 4c, d).

To estimate the single capillary occlusion resulted in the changes of vascular architecture and hemodynamics of connected capillary networks, we documented the velocity and diameter changes generated by occlusion in layer 2/3 of the somatosensory cortex of 161 vessels in 13 mice (Supplementary Fig. 6a). We found that there was no significant difference in the corresponding relationship between diameter and RBC speed of all vessels before and after occlusion, but reduction in blood flow appeared after occlusion (Supplementary Fig. 6b). Averaged RBC speed was significantly lower of total capillaries (1.234 ± 1.075 mm/s for before vs. 0.980± 1.147 mm/s for after; p = 0.0068), while the average diameter of total capillaries showed a small but significant increase after occlusion (4.693 ± 1.166μm for before, 4.911 ± 1.220 μm for after; p < 0.0001) (Supplementary Fig. 6c). However, the upstream and downstream vessels showed completely different hemodynamics after micro-occlusion. A complicated pattern of reversed and non-reversed flow is observed in vessels farther from the clotted capillary, especially in downstream vessels (Fig. 4f, Supplementary Fig. 6a). The ratio of RBCs speed and diameter to baseline of each capillary was regarded as a reference to evaluate the changes in hemodynamics and vascular diameter at various orders after occlusion (Fig. 4f, g). For all upstream and side branch vessels, the average blood velocity was increased by 15.3 ± 72.4% to baseline, and slightly increased 7.3 ± 11.8% (±SD) for diameter (Fig. 4f, g). The RBC speed of one to three upstream branches (U1, U2, U3) changed dramatically, with flow speed increase of 32 ± 58.5%, 13.4 ± 52.1% and decreased of 34.5 ± 80.5% from U3 to U1 (for U1 and U2, p = 0.0218; for U1 and U3, p = 0.0446;) (Fig. 4f). Interestingly, we observed complex changes in U1 after occlusion, whereas stall and reversal flow occasionally appeared (Fig. 4f, Supplementary Fig. 8a), in which the reversal flow came from TUS, originally from the neighboring diving artery-supported vascular network. On average, all upstream branches had slight dilation, among which U2 dilated most for 11.5 ± 10%, while U3 had 8.1 ± 8% and U1 had only 1.7 ± 8% (for U1 and U2, p = 0.0103;) (Fig. 4g). Upstream and upstream side branches showed significant different in RBC speed (0.970 ± 0.720 for upstream, 1.422 ± 0.654 for side branches; p = 0.0154,), while no difference in the diameter changes after occlusion (Fig. 4h), which may due to compensation of the increase flow for the flow deficit in target vessel. The downstream vessels from the clot showed a diversity of hemodynamic changes, on average, in which the flow velocity reduced to 36.3 ± 72% in D1 and 59.2 ± 100.1% in D2, while D3 reduced to 96.1 ± 91.6% (for D1 and D3, p = 0.0112;) (Fig. 4i). There occasionally occurred reversal flow in some downstream capillaries after occlusion, which may be induced by the deficit perfusion source of the clotted capillary and rescued by the adjacent vessel network supported by another diving artery. In addition, the diameter had significant difference in D1 to D3: 98.5 ± 8.2% for D1; 102.2 ± 7.8% for D2; 105.3 ± 8.0% for D3 (for D1nd D3, p = 0.0153) (Fig. 4f).

In general, the capillary network in layer 2/3 had significant difference and functional diversity of flow velocity and diameter change between various orders of branches after occlusion. On average, the upstream vessels showed strong vasodilation and increased flow velocity, especially in upstream side branches, while the downstream vessels showed dramatic decreased blood flow, and sundry but no significant change in diameter. This finding suggested that acute capillary dilation in upstream vessels contributed in robust redistribution to adjacent tissues, and the clotted capillary led to flow resistance that progressively worsened restrict microcirculation within the ischemia downstream basin, while downstream vessels reversed blood flow sometimes to achieve redistribution of blood flow and new pathway of perfusion.

In addition, we measured the evolution of these capillaries over a period of 1 hour, acquired the morphology and patency of individual capillary segments of baseline and 5, 15, 30, 45, 60 min post-occlusion (Supplementary Fig. 9). Each capillary segment was categorized based on the forms of pathology exhibited, including (1) constant (blood keeping flow), (2) reverse, (3) stall, (4) BBB leakage [34]. In these analyses, we did not distinguish between different types of obstructions, such as clot, RBC or leukocyte, as all could have contributed to impairment of flow. Blood-Brain-Barrier break down was identified when intravascular FITC-dextran dye had extravasated into the surrounding tissue. When capillary responses were examined on a vessel-by-vessel basis over the imaged period, we observed substantial heterogeneity in both the timing and duration of constant and reverse events. To understand the overall dynamics of capillary over time, we considered the capillaries as a population of individual mouse. An average of 15% of the total capillaries measured lost flow at 15min after occlusion, as assessed by the cessation of movement of RBCs within the lumen (Supplementary Fig. 7; blue line). Part of the stagnant capillaries restored flow gradually, which resulted in the proportion of the constant vessel raised from ~82% at 15 minutes to 86% at 1 hour (Supplementary Fig. 7; black line). The decreased proportion of stalled vessels was partly caused by BBB leakage. The degradation of the BBB and leakage of circulating FITC-dextran into the parenchyma was a comparatively late event, showing at 15 minutes post-occlusion and affecting ~3% of the total microvessels at 1 hour (Supplementary Fig. 7; green line). The capillary dysfunction observed *in vivo* preceded and likely contributed to the expansion of the occlusion at capillary level did not lead to a complete cessation of flow but to a redistribution of flow.

### Micro-occlusion induced layer specific neurodegeneration

The stable blood outflow of capillaries contributes to the robustness of perfusion, which is crucial to guarantee a sustained nutrient and O^2^ supply throughout the tissue. Dendrite and spine structure are relatively resistant to moderate ischemia, but large scale of ischemia leads to stroke, eventually leading neuron death and cause the neural circuits remodeling within several hours to months [35–37]. Even, neurons that are deprived of their normal substrates can show signs of widespread loss of dendritic structure after as little as few min of ischemia then delayed cell death [38]. But relatively little is known about the precise spatial relationship between capillaries blood flow and neurons states, especially the spatial and temporal pattern of micro-ischemic damage. To explore the single or several capillaries occlusion effects on the neural structure change, we produce micro-occlusion of capillaries surrounding a GFP labelled neuron soma in 2/3 layer of somatosensory cortex and investigated the integrity of neuron structure immediately (Fig. 5a, b).

**Figure 5.**
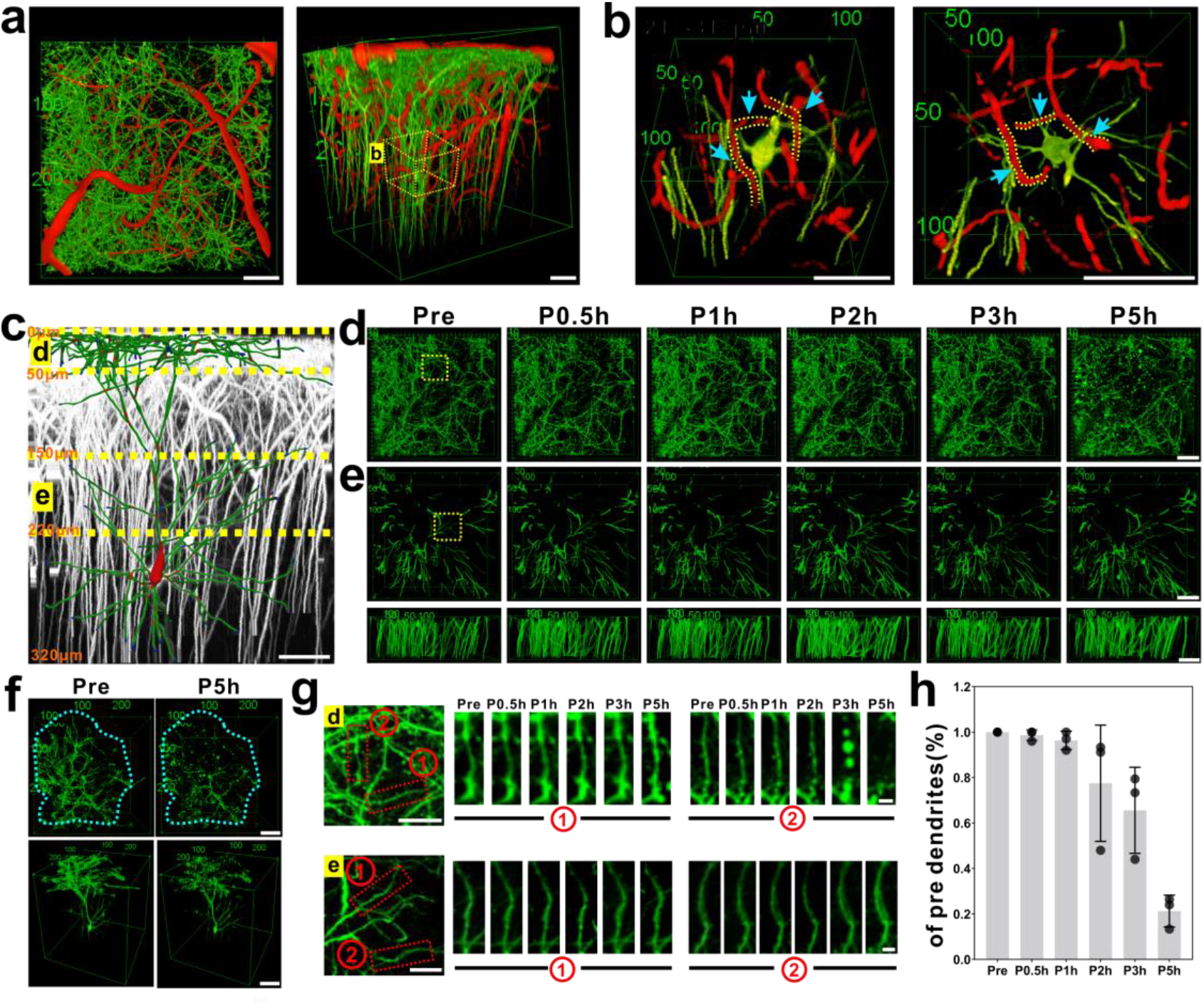
Localized small strokes of several capillaries near to the soma escape immediate laminar distribution of damage to dendrites a. 3D TPM reconstructions of maximal intensity projection of 0-332μm of the cortex indicating the vasculature and neuron axon and dendrites. (Left: top view; right: side view. Red: TRITC-dextran; green: gfp) scale bar, 50μm. b. Enlarged 3D reconstruction of the targeted soma and nearby capillaries form planar a (yellow dashed box), showing the vasculature and soma position before photothrombosis. (Left: side view; right: top view. Blue arrow and yellow dashed line refer to the capillaries clotted by photothrombosis) scale bar, 50μm. c. Side view of 3D rending imaging of the axons and dendrites (white). Target neuron was distinguished by color assignment (red: soma and nodes; green: axons and dendrites). Scale bar, 50μm. d. Top view of neuron axon and dendrites during different timeline before and after photothrombosis in 0-50 μm. Scale bar, 50μm. e. Top view of neuron axon and dendrites during different timeline before and after photothrombosis in 150-220μm. Downside: Side view of the same area. Scale bar, 50μm. f. Top view of the whole neuron axon and dendrites structure before and after photothrombosis. Blue circle represents the main area of the structure. Downside: 3D reconstruction of the targeted structure. g. Magnified view (red box in planar e) showing dendrite structural changes wthin 5 hours. 1 refer the dendrite from the target neuron, 2 refer the dendrite from the non-target neuron. e. Dendritic structure (a sub-stack Z-projection, 0-100μm from surface) pre and 0.5h, 1h, 2h, 3h and 5h after photothrombosis showing a stable dendrite (down) and a lost dendrite (up) h. Dendrite number expressed as a percentage of the number present after photothrombosis in each mouse.

Firstly, we blocked single capillary near the target neuron, mapped the spatiotemporal relationship between acute capillaries ischemia and apical and basal dendritic tufts structure within single neurons *in vivo* over 5 hours. We found that there was no blebbing or visible damage to all tuft spiny dendritic structures of the targeted neurons, which proved that the ischemia of a single capillary had limited impact on the nearby neuron, and the neuron may be supported by other surrounding capillaries (Supplementary Fig. 8). As previous work had indicated that the mean distance between the neurons to the closest micro vessel was 17.8 µm [31, 32], we blocked all surrounding capillaries near by the targeted soma within 50μm. Specifically, we investigated how the occlusion of all surrounding capillaries led to propagating damage throughout axon and dendrites of the neuron soma located in the ischemia center. Dendritic tufts are the dendritic region where are most vulnerable to ischemic damage. Focal bead-like swelling in dendrites and axons is thought to be a neuropathological sign of ischemia associated with neuronal dysfunction and precede cell death [39, 40]. We analyzed large apical and basal dendrites and their tuft extensions to quantify the apparent laminar distribution of ischemic damage to neurons. The entire dendritic structure from the pial surface to the cell body were reconstructed (Fig5c, f). In three animals after micro-occlusion, we observed intense blebbing within the apical dendritic tuft and a complete loss of dendritic spines in this region which belongs to the targeted neuron in layer 1 over 5 hours (Fig. 5d). Ischemic damage to the targeted neurons located at the ischemia core was contained within apical tuft spiny dendritic structures and did not propagate to spines on the more proximal region of the soma (Fig. 5f, g). Dendrite counting indicated that over a 5–6 h period post micron-ischemia, the total number of dendrites was gradually reduced in layer 1, suggesting some degree of progressive fine structural damage (Fig. 5h). However, we rarely observed the blistering of dendrites and loss of spines on the dendritic tufts of non-target neurons (Supplementary Fig. 9). Moving down the dendritic tree, the remaining portion of the basal dendrite extending 150 to 220μm below the surface of the cortex appeared to be intact as indicated by the presence of spines within these regions (Fig. 5e). To verify that other neurons located nearby the micro-occlusion core could be free of ischemic damage, we also analyzed the neurons located within 100μm from the ischemic core over 5 hours, and found no visible damage to all tuft spiny dendritic structures (Supplementary Fig. 9). This analysis revealed that the spines and dendritic structure of these neurons could be intact throughout all cortical layers, including the most superficial dendritic tuft. All these data verified that micro-occlusion in a restrict area had a very limited effect on peripheral neurons. In other words, the complete breakdown of blood flow in a restrict area could cause a continuous devastating blow to the inside neurons in several hours, but not to neighboring neurons.

## Discussion

We established a highly reproducible robust *in vivo* model of cortical vascular photothrombosis, compared with conventional approaches, which allows faster, more precise and controlled clotting of single/several vessels in depth. Two photon absorption–based photothrombosis prevented damages to other neighbor parenchyma or vessels. This allows us to produce multisets capillary occlusion within seconds to mimetic the small vessel diseases. We found that micro-occlusion at layer 2/3 and layer 4 within the same vascular arbor resulted in distinct effects on downstream flow redistribution. Analysis of the microvascular architecture and blood-flow dynamics after single capillary occlusion reveals distinct autoregulation in upstream vs downstream. Further, micro-occlusion of capillaries surrounding a labelled target neuron induced dramatic and rapid changes in neuronal dendritic architecture. In sum, our technique enhances the precision and power of photothrombotic study. Which enable the creation of robust models of any type of occlusion, combined with the ability of multi-photon imaging to measure cerebral blood flow during normal and pathologic states is critical for the development of an animal model of neurovascular disease.

### The mechanism of ultra-fast light induced photothrombosis

In an attempt to provide means of altering cerebrovascular dynamics in a controllable fashion, photothrombotic models, using Rose Bengal (RB) or other agents, have been developed, which exploit reactive oxygen species generated from focused light, causing inflammation and damage to cellular vascular walls resulting in microinfarcts. Rose Bengal is the most efficient known photodynamic generator of singlet molecular oxygen to induce platelet aggregation during photothrombosis using the 532nm laser [17]. One of the major challenges of RB photothrombosis relates to out of focus excitation when exposing the brain to light, as it could lead to undesired blockage of adjacent vessels precluding the blockage of a single capillary, although which is currently mitigated through various strategies. Nevertheless, these solutions nonetheless still create non desired excitation in the vicinity of the targeted vessel by diffused light and is limited in penetration depth due to the short wavelength of the excitation light used to create the blood clot. Alternative mechanisms to create microinfarcts have also been proposed, relying solely on energy deposition from near-infrared ultrafast laser pulses to cause localized burning injury-inducing parenchymal microvessel damage while preserving the overlying vasculature developed for several years to research the superficial vessel functions [22, 30, 41].

Here we proposed a novel model of highly targeted photothrombosis, through multiphotonic excitation of RB. In this work, clot formation from the RB excitation was used as an indicator of oxidative stress from the RB mechanism of photothrombosis. We used spiral scan to induce photothrombosis which led to high efficacy of volumes’ excitation creating a bigger number of oxygen singlets, and triggered the endogenous clotting cascade. One could verify with sophisticated probes if the singlet oxygen agents are emitted by RB. In this trial, some bigger vessels were also targeted (30-50 um), resulted in a vascular insult after much longer photo irradiation, which could testify the efficient oxygen singlet creation via multiphoton activation of RB. Moreover, trials were performed on mice models and shows that a success rate of above 95% is obtainable with RB excitation at 1070nm.

### Layer-specific microvascular network redistribution

The mammalian neocortex is organized as a six layer with different neuron types and density structure as a functional unit. The microvascular architecture in the cortex also form highly interconnected loops in six layers to support the neuron function, which shows a broad variation in the density of the vasculature as a function of depth in mouse, monkey and human [2, 27, 42, 43]. The neurovascular mechanism of how the microvascular be modulated to support the neuron activity is the leading question in the field. Previous data with anatomy analysis of the fine structure of the microvessels relationship with the neuron structure revealed vascular topology structure [2, 32]. Using the ultrashort laser pulses to specifically occlude the single penetration vessels, the patterns of blood flow showed that microvessels do not provide enough collateral flow to perfuse tissue which indicate that the penetrating vessels was the bottleneck in the cortex[25, 41, 44]. Computational analysis of microcirculatory blood flow and pressure drops further indicates that the capillary bed in deep layers, including capillaries adjacent to feeding arterioles (d<6 mm), are the largest contributors to hydraulic resistance [45]. However, experimental evidence has been able to identify high-branching-order capillaries release O_2_ acting as reserve that is recruited during increased neuronal activity with pO_2_ tracer under 2-photon microscopy [46–49].

We found that the capillary network in layer 4 showed a higher autonomic regulation ability than that in layer 2/3 responding to the acute occlusion of the pre-capillary. This finding reflects the layer-specific differences in vascular density and pressure drop characteristics. Right after acute occlusion, the capillary network had some determine mechanisms to cope with the drastic decrease in flow source, which was also related to the surrounding DA.

### Local capillary network flow autoregulation

Changes in capillary morphology as well as evaluation of dynamic flow patterns plays an important role in microcirculatory dysfunction [50–53]. Even though the structure of the capillary bed is commonly described as being homogeneous its flow field is highly heterogeneous. The robustness of the capillary bed can be explained by its mesh-like structure, which is beneficial for an efficient redistribution of flow. Capillary dysfunction ends up with a diminished functional hemodynamic response and oxygen extraction [53]. The capillary bed is a highly interconnected network that allows blood flow to efficiently re-route around local vascular defects. A logical hypothesis would be the continued or increased flow of neighboring capillaries and a compensation for perfusion deficits incurred by the occluded capillary. Upstream capillaries should therefore also remain patent to serve the efflux.

Our data showed the robust redistribution of the upstream and downstream vessel blood flow. Similar to the previous pial and penetrating vessels results, occlusion experiments have been performed to assess the overall robustness of the capillary bed in the distribution of flow [30, 44]. Previous results showed that the impact of a microvessel occlusion is minimal and that the RBC flux recovered to 45% of its baseline value by three branches downstream of the occlusion.

Furthermore, Other data[54] show that 63% of all capillaries are fed by more than one DA and hence a redundancy towards DA occlusion persists as well. These findings may contribute to optimization of microcirculatory flow patterns in microcirculatory dysfunction disease.

### Micro-ischemia related to neurodegeneration

Accumulating evidence suggests that cerebrovascular changes can occur prior to the presentation of symptoms and promote many neurodegeneration [55–58]. Large-scale stroke models, such as distal MCAO, and photothermolysis with RB with 540nm green light induced mini-stroke have been widely used in mouse model to reveal the stoke pathology after ischemia for a long time [59–62]. These methods induced brain tissue ischemia have proofed that many neurodegeneration diseases such as AD and ALS involved in the subsequent effects [63–66]. But how the capillary system induced ischemia correlated with the neurodegeneration is still lacking pathological evidences. Recent results showed that in mouse and humans with cognitive decline, amyloid β (Aβ) constricts brain capillaries at pericyte locations, inhibiting the capillary constriction caused by Aβ could potentially reduce energy lack and neurodegeneration in AD [67].. Our method provides new tools to investigate the neuron-astrocyte-capillary interaction in the single capillary level. Micro-occlusion in capillary also induced BBB disruption, which allows influx into the brain of neurotoxic blood-derived debris, cells and microbial pathogens and is associated with inflammatory and immune responses, which can initiate multiple pathways of neurodegeneration [68]. Also, the implication of the microvascular component has been suggested for several other disorders affecting higher cognitive functions such as ASD [69, 70], schizophrenia[71], or obesity [72].

The limit of oxygen diffusion is 100–150 μm in live tissue. The distance between capillaries is ~40 μm in mice[73]. The distance over which oxygen can diffuse in the normal brain is highly regulated. Vessel dysfunction especially occlusion leads to a decrease in blood-vessel density and causes the inter-capillary distance to exceed the limit of oxygen diffusion, and the consequent reduction of oxygen availability to brain cells may result in stress on neurons or glial cells during periods of increased activity in a particular brain region. Any dramatic alterations (remodeling, regression, angiogenesis, etc.) in the pattern of the brain vasculature may result in altered regulation of brain microcirculation, which in turn is likely to lead to changes in the local supply of glucose and oxygen in the brain [74–76]. Neurons are the key player in brain function, and they consume the most glucose and oxygen in the adult mammalian brain[77]. After an alteration to the microcirculation, especially a decrease in blood supply followed by vessel regression, neurons will be the most vulnerable cell type under these stressed conditions, and they likely to sustain damage [78]. We wished to uncover whether this micro-ischemia damage would propagate into the farther dendritic and axonal compartments as well as what the spatial and temporal pattern of this ischemic damage would be. And examination of the integrity of tissue immediately adjacent to the ischemia core is important as this is where changes such as adaptive neuronal plasticity [79] and circuit reorganization[80] that underlie functional recovery are thought to occur. Although we detected a substantial damage of neuronal structure in the cerebral cortex due to the targeted capillaries occlusion, further work is required to determine how blood vessel occlusion regulates energy-related metabolism under physiological conditions and whether a certain threshold of vessel occlusion and regression leads to neuronal or synaptic degeneration. Our approach allows the investigation of the disruption of neurovascular units *in vivo* under ischemic stroke in a precise manner, which could also help investigate the relationship between neuronal networks dysfunctions and metabolism in the capillary level combining with the multiphoton imaging microscopy. Our technique provides a new powerful tool of investigation by making micro-scale function changes of microvascular topology in disease models accessible.

The present study complements past work that focused on large ischemic and hemorrhagic disruptions, which helped to define the molecular basis for neurovascular pathology. Diseases of small vessels are common causes of progressive disability and cognitive decline [50, 81]. Our induction of microvessel dysfunction may provide a unique means to model micro-occlusion, that is, localized, small ischemia infracts observed in aged human patients [82]. Postmortem results from the brains of patients with dementia have shown many microinfarcts with diameters of 0.1 to 1 mm [30, 83, 84] . We can manipulate various types, depths and sizes of vascular occlusion to obtain different infarct pathologies, which facilitates a microinfarct strategy to study brain injury and repair coupled with live-imaging technology. Our methodology provides a useful tool to perturb and measure coupling, a long-standing challenge in neurovascular research[85].

In addition, this is a precise microsurgery type anti-vascular method and holds particular promise for treating diseases in brain cortex, where high spatial selectivity is critical for preventing collateral effects on central nervous system function. It represents a precision medicine approach for vascular disease treatment on a per-vessel/per-lesion basis.

## Methods

### Animal and surgery

All experimental procedures were approved by the Institutional Animal Care and Use Committee of Zhejiang University and in accordance with the Institutes of Health Guide for the Care and Use of Laboratory Animals. Male C57BL/6J mice (9-10 weeks old) and Thy1-EGFP mice (9-10 weeks old, male and female) were housed at 24°C with a normal 12 h light/dark cycle, and fed with water and food.

Mice were anesthetized with isoflurane (5% inhalation, mixed with pure O2, 0.5L/min) and placed in a stereotactic frame, after which isoflurane was maintained at 1%–2% throughout the procedure. Body temperature was maintained at 37°C with a heating pad. The craniotomy procedure was performed as previously described [86]. A craniotomy (measuring 4 mm in diameter) whose focus was at 0.3mm anterior, 2.3mm lateral to bregma was exposed in the right sensory cortex using a dental drill. The dura was meticulously moved and artificial cerebrospinal fluid (ACSF) filled the exposed cortex. Lastly, the round coverslip (diameter: 8mm, and thickness: 0.17mm) was used to cover the brain tissue, and medical glue was used to seal its edge. Dental cement was subsequently used to fortify a custom-made stainless-steel headpost with screw holes, allowing the mice to be head-fixed while imaged. Mice received a subcutaneous injection of buprenorphine (0.05 mg/kg) for at least three days post-surgery and were allowed to recover for at least two weeks before they were imaged.

### Two-photon Imaging

All animals with a craniotomy window were mounted and immobilized to a stationary apparatus with a heatpad and were imaged by TPM in an anesthetized state (1-3% isoflurane). 20 minutes before photothrombosis mice were retro-orbitally injected with 0.05 mL 2MDa TRITC-dextran (10mg/mL, Aladdin) or 0.05 mL 2 MDa Fitc-dextran (10mg/mL; Sigma) to provide a fluorescent angiogram *in vivo* in different conditions. For Neural (GFP) and vascular (TRITC) signals, dual-channel were recorded by a customized TPM (Ultima IV, Bruker Corporation, 500-550 nm filter (green), and 570-620 nm filter (red)) coupled with a Femto Second laser (Chameleon Ultra II, Coherent Inc. model-locked Ti: Sapphire laser) which generated two-photon excitation at 860 nm, equipped with a 16x (numerical aperture = 0.8) water-immersion objective (Olympus). The two-photon–excited fluorescence is reflected by a dichroic mirror and relayed to photomultiplier tube (PMT). PMT settings and laser power was adjusted by Pockels Cell (Conoptics) according to experimental requirements. For planar structure imaging, the frame FOV was 375.47um×375.47um (pixel: 1024 × 1024) with ~1Hz frame rate. For z-scan imaging, the same size of FOV but different pixels (512 × 512) were chosen with ~2Hz frame rate, 1μm of z-axis step. Each frame was repeated four times, and the averaged frame was taken as the result. For RBC velocity measurement in real-time, we used line-scan measurement, repetitive line scans along the axis of the vessel, at a rate of at least 800Hz, were used to form a space-time image in which moving RBCs produce streaks with a slope that is equal to the inverse of the speed (Supplementary Fig. 2). And we typically acquired line-scans for at least 10s totally for one vessel and report the average speed over this period. At the same time, it is also to eliminate the influence of blood flow changes caused by the regular dilation and contraction of the blood vessels in the resting state under anesthesia. Considering that multiple and long-term collections are used for average speed and the complicated blood flow of each blood vessel, we do not consider the change of blood flow when calculating blood flow speed.

In mouse, Rose Bengal is cleared from circulation within 5 min[60], we typically performed laser stimulation within 5 min of intravenous injection of Rose Bengal. Generally, the structure and blood flow velocity after acute occlusion are collected 15 minutes later. For the RBC speed measurement of the capillary network within 1 hour, measured at 10 minutes before occlusion 15 after photothrombosis.

### Targeted vessels photothrombosis

Once the baseline measurements of vessel structure and RBC speed were completed, immediately followed a retro-orbital vein injection of ~50 μL of Rose Bengal, prepared at 1.25% (w/v) in sterile saline. Irradiation of the Rose Bengal leads to the production of singlet oxygen, which damages the wall of the vessel and subsequently triggers a clotting cascade that leads to an occlusion [17, 18, 87]. A Femtosecond 1070nm uncaging laser (Fidelity 2, Coherent) was used as the irradiation light which directed onto the microscope through a galvo-galvo system whose power regulated by the Pockels Cells (Conoptics) (Fig. 1). We formulated a series of parameters such as duration, diameter and laser power etc. to deliver the laser stimulation pattern to the target vessel by spiral scans (Fig. 1). When marking a spiral scan, the laser starts at the center and, travelling at a constant ultra-fast speed over the course of the chosen duration, spirals outward to the edge circling about the center for the number of revolutions specified. The size of the spiral (same as the diameter of the targeted vessel), the number of revolutions (how many revolutions are made to make it from the center of the spiral to the edge, usually 2 times to the diameter of targeted vessel), stimulation duration (3s in our paradigm), step size (~670 nm) and repetition were specified on the vessel types and sizes (table 1). The laser power was controlled from 122mW to 158mW under the objective altering to the depth of the target blood vessel. The paradigm of light stimulation is 3s stimulation following 2s imaging when we can monitor the real-time changes of blood vessels, which were repeated 3-60 times (15-300s) mainly according to the vessel type and depth. (table1) Once the clot formatted, the stimulation paradigm stopped immediately.

### Image analysis

All data analyses were performed using MATLAB (R2020b; MathWorks), iMaris (Bitplane), neuTube [88], and ImageJ (ImageJ, US). The reconstruction and rendering of 3D images used the 3D viewer plugin in ImageJ; in addition, neuTube is used for neuronal structure, vascular network tracking, and the RBC speed was colorized using the MATLAB code customized by the laboratory. For diameter measurement, average width across the length of the selected segment was calculated as the vessel diameter. Capillary diameters were calculated as the FWHM of the intensity profile across the capillary width [89]. For RBC velocity measurement, LS-PIV in MATLAB was used (Rong A. Wang et al, https://sourceforge.net/projects/lspivsupplement/files/). The slope was calculated using an automated image-processing algorithm. Measured parameters were then mapped to vascular networks that included the targeted vessel. For motion correction, images were processed with Image Stabilizer in ImageJ (Kang Li, Steven Kang, http://www.cs.cmu.edu/~kangli/code/Image_Stabilizer.html).

### Statistical Analysis

All statistical analyses were performed using GraphPad Prism (GraphPad Software, San Diego, CA, USA) software. Statistical analyses were performed using unpaired, two-tailed t-test and ANOVA test. Analysis of rating scores was performed with a non-parametric Mann-Whitney U-test or Kruskal-Wallis test. P < 0.05 was accepted as statistically significant. Results were expressed as mean ± standard deviation. *P < 0.05, **P < 0.01, ***P < 0.001 and ****P < 0.0001.

## Acknowledgments

This work was supported by the National Natural Science Foundation of China (91632105, 81630098), the Zhejiang Provincial Natural Science Foundation of China (LY17C090005), and the Fundamental Research Funds for the Central Universities (2019QNA5001).

