## Supplementary figures and images for "Imaging microvasculature network evolution and neurodegeneration with precise photothrombosis approach"

### Supplementary Fig.1

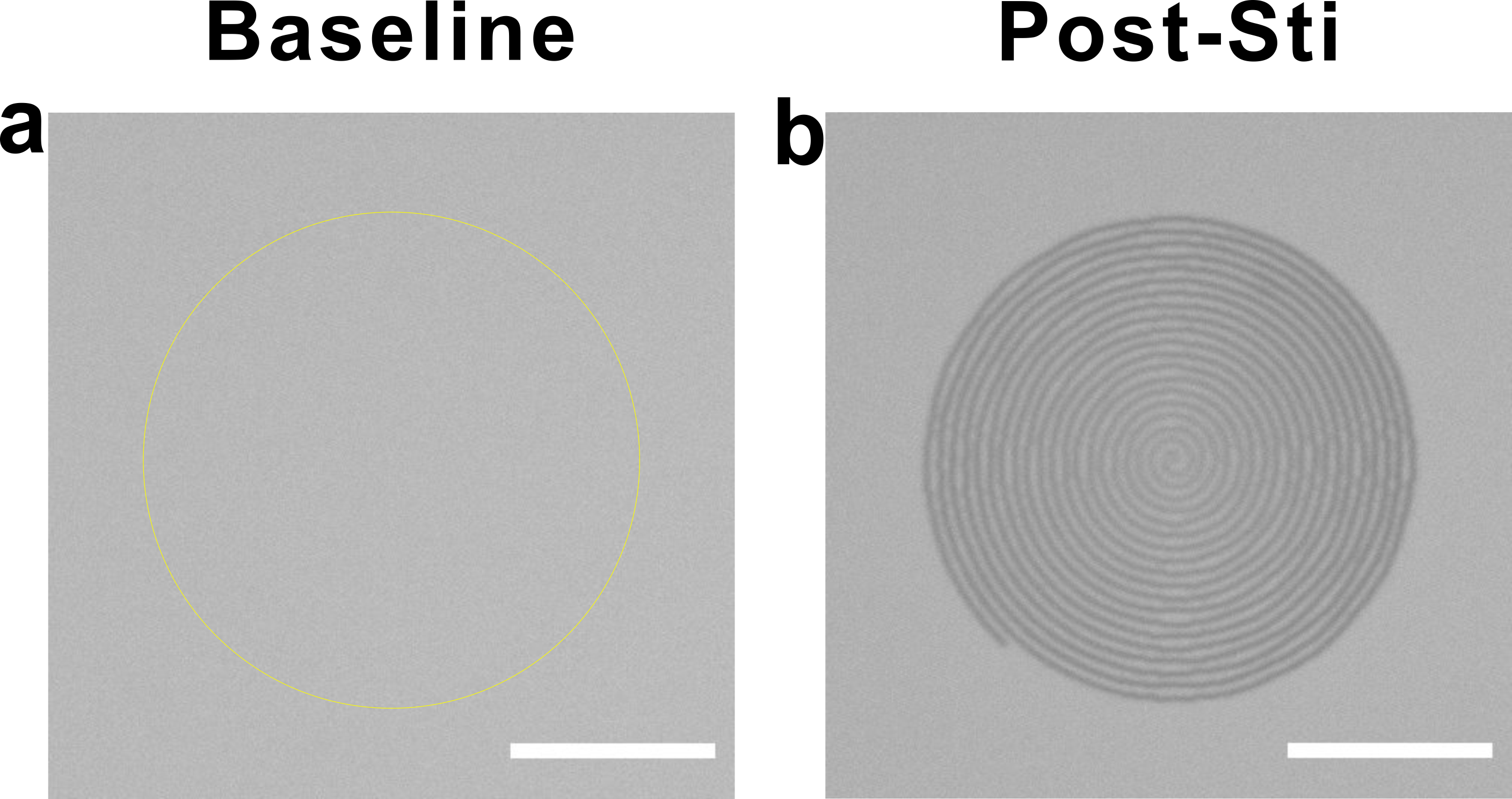

### Supplementary Fig.2

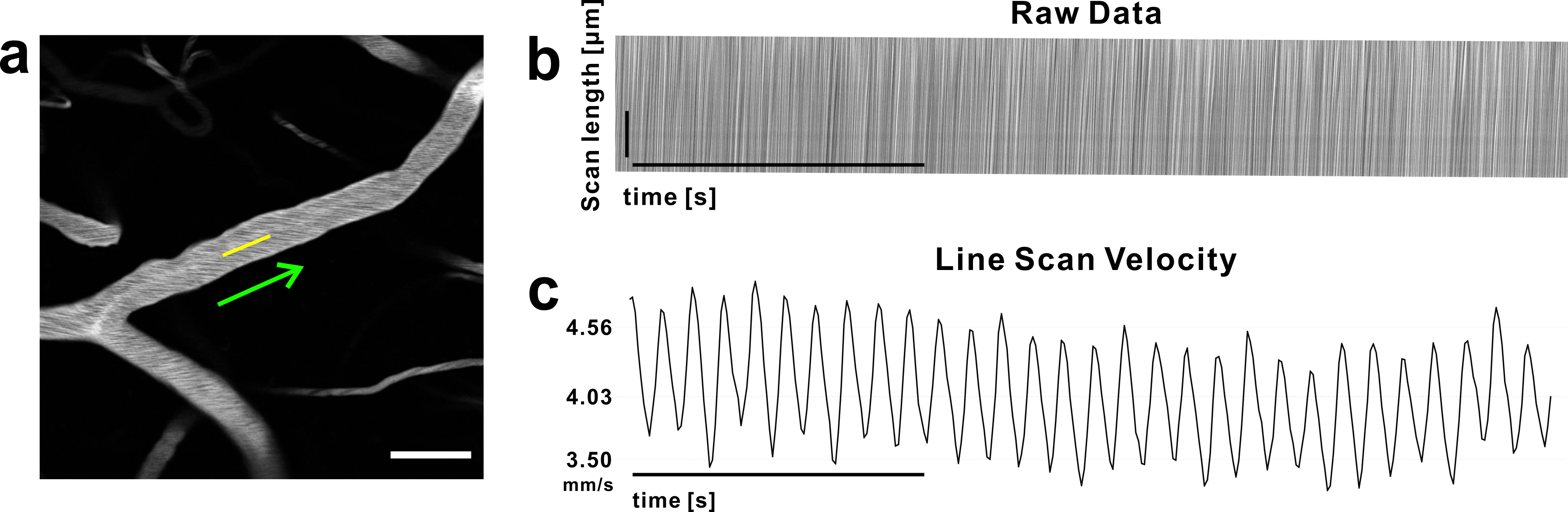

### Supplementary Fig.3

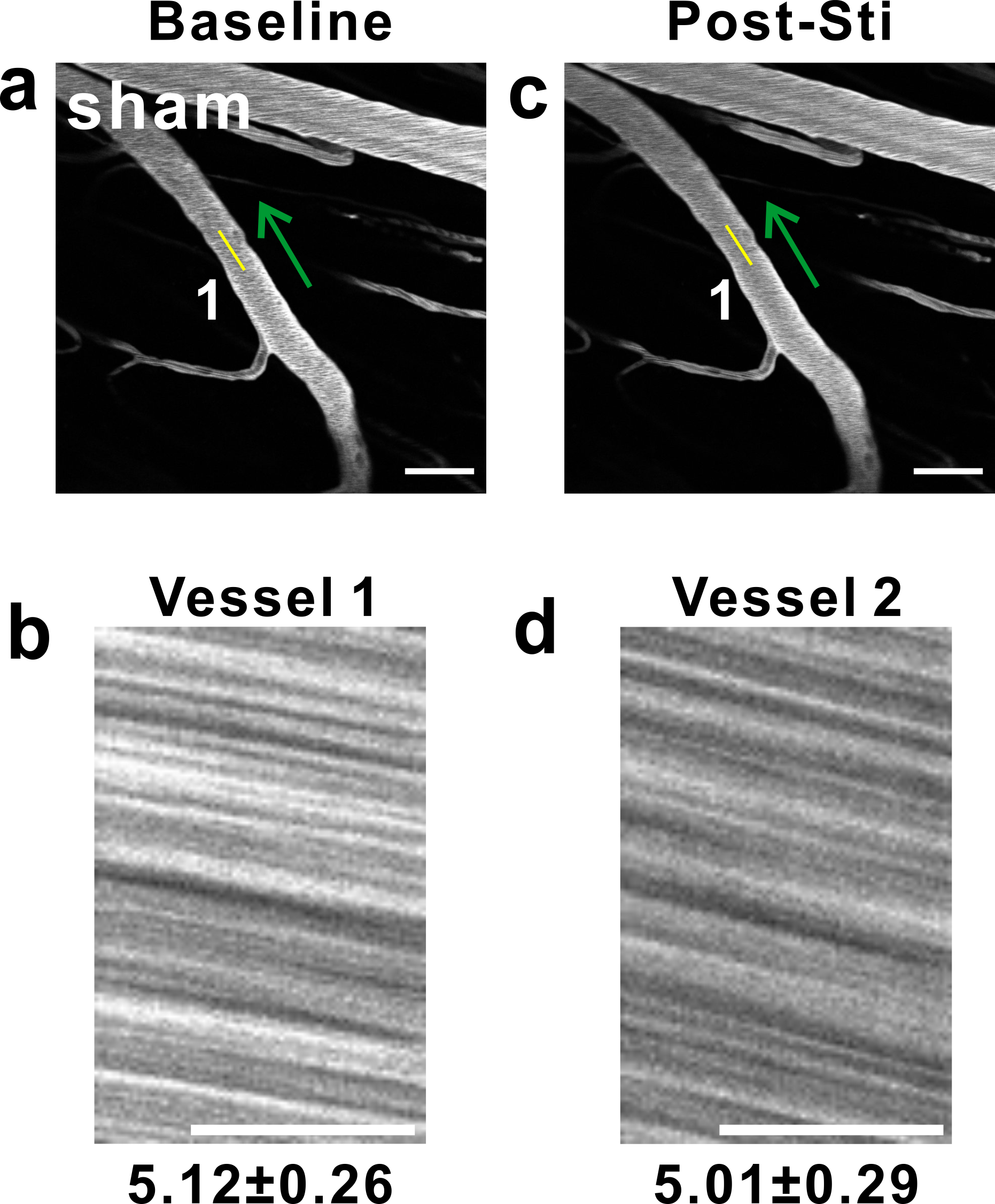

### Supplementary Fig.4

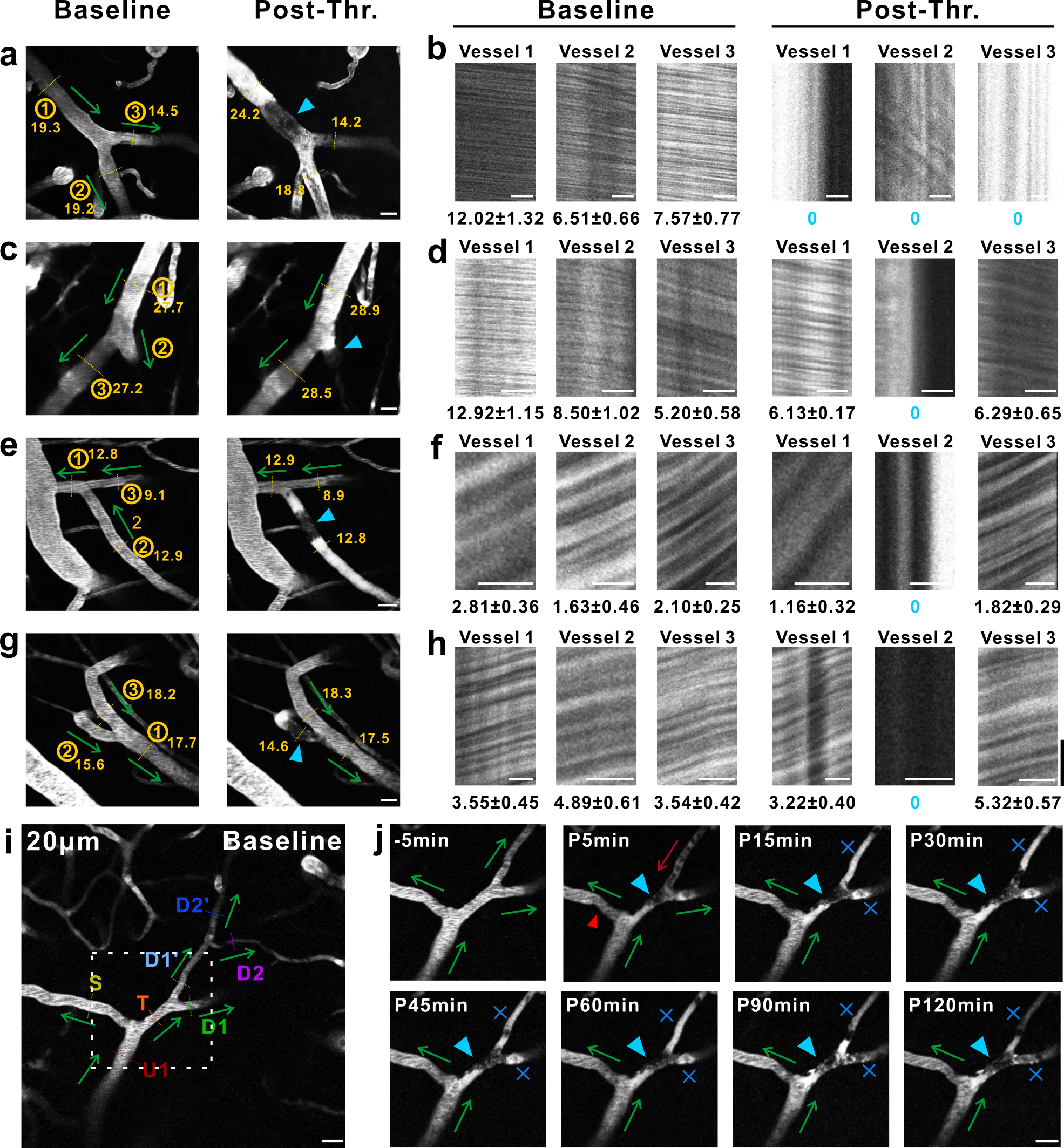

### Supplementary Fig.5

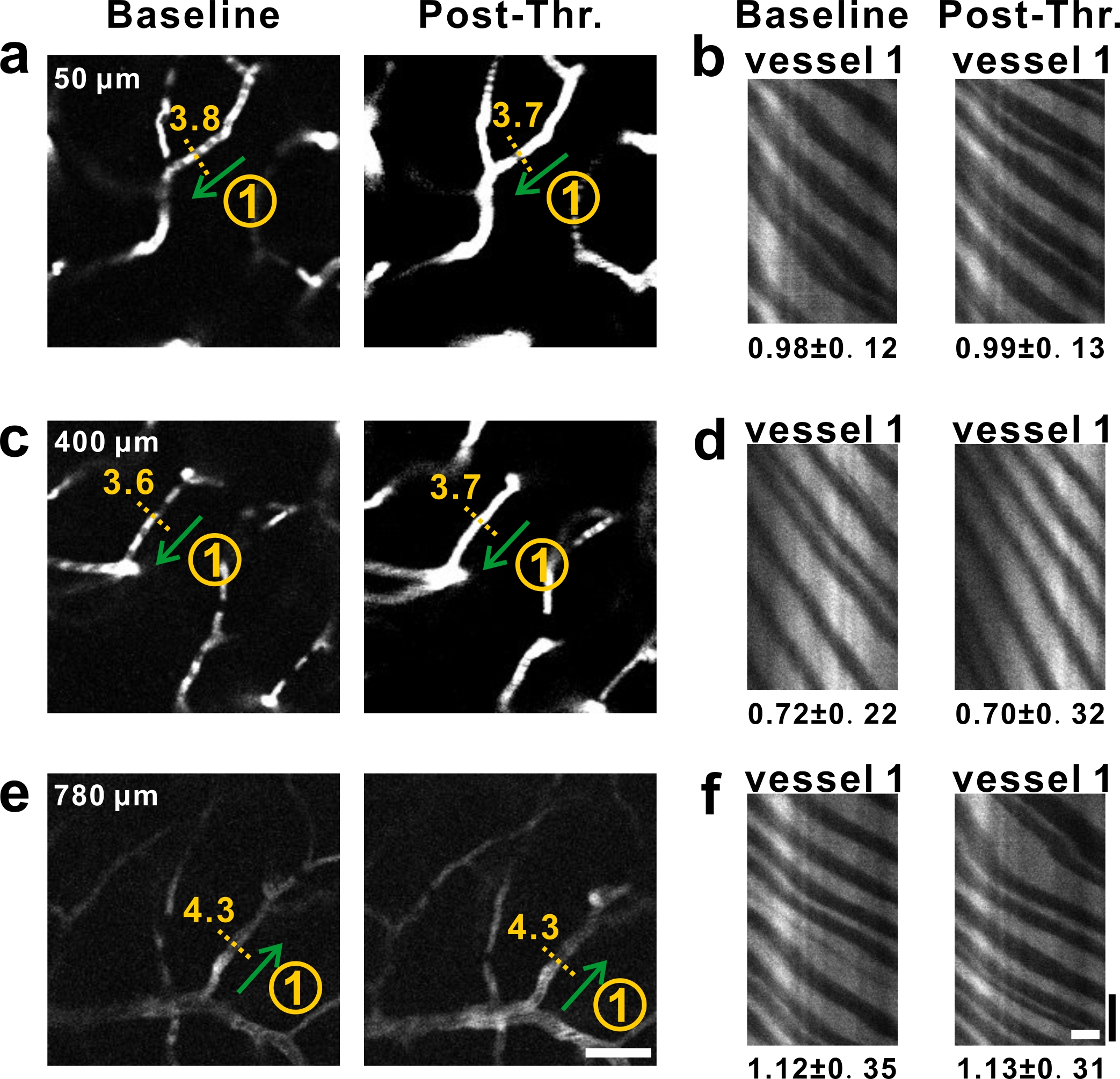

### Supplementary Fig.6

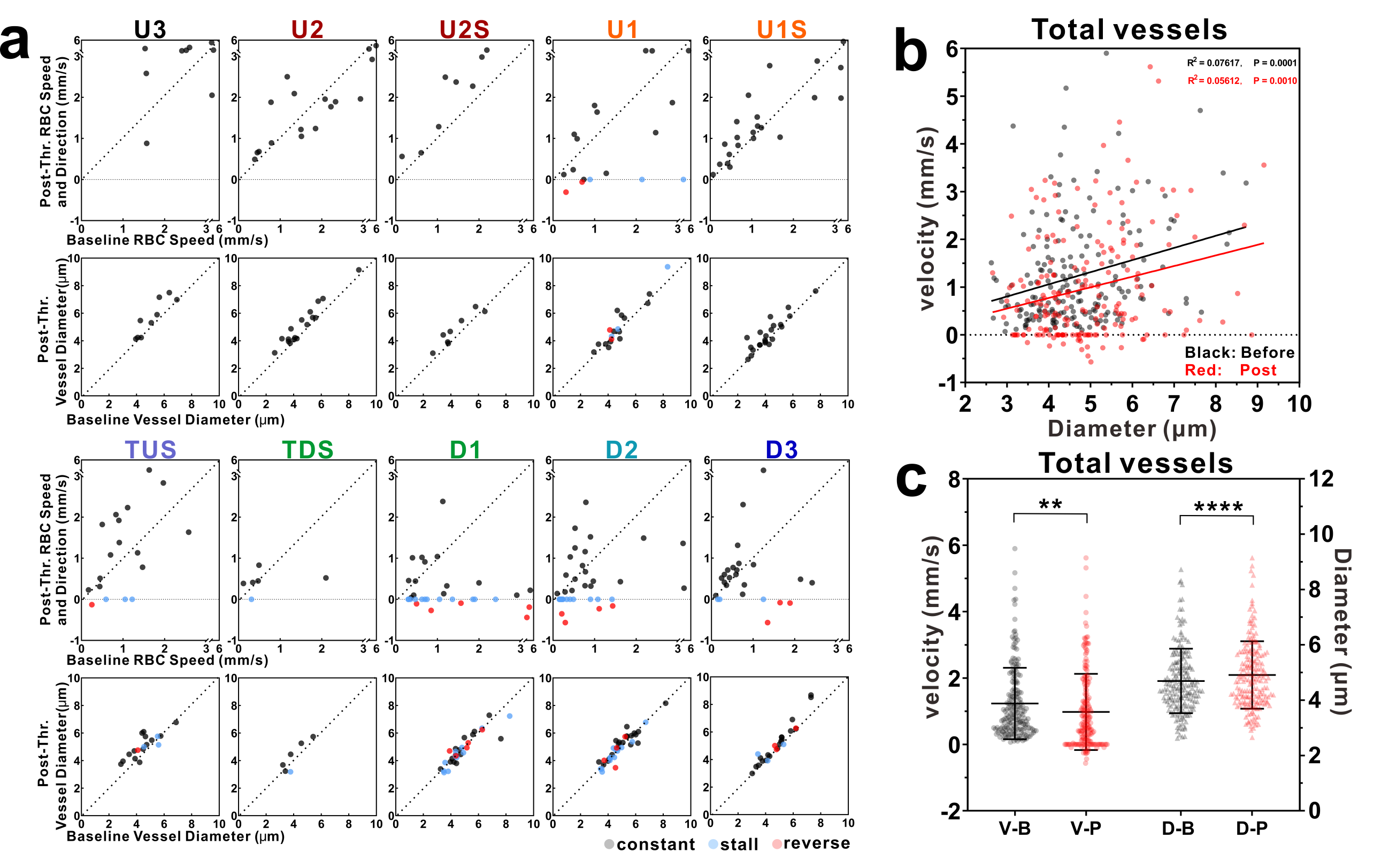

### Supplementary Fig.7

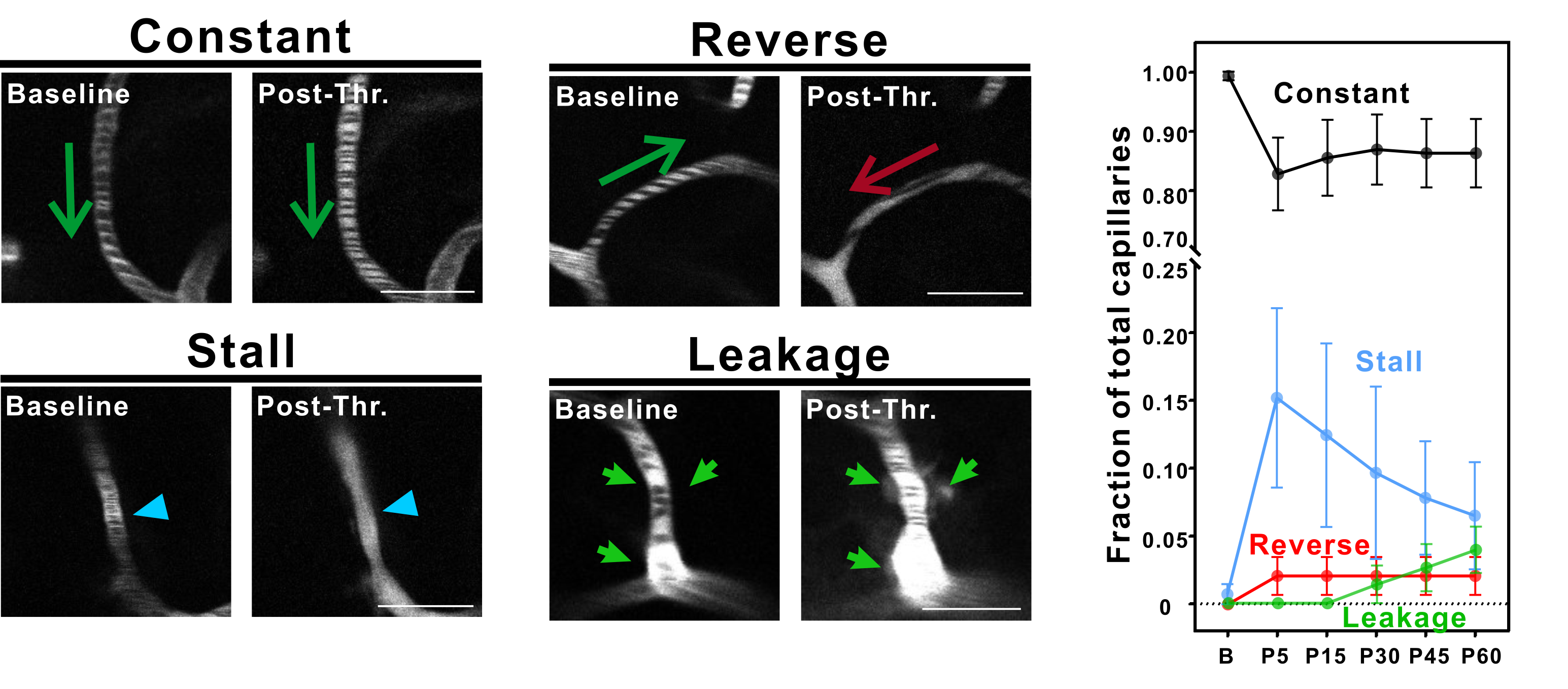

### Supplementary Fig.8

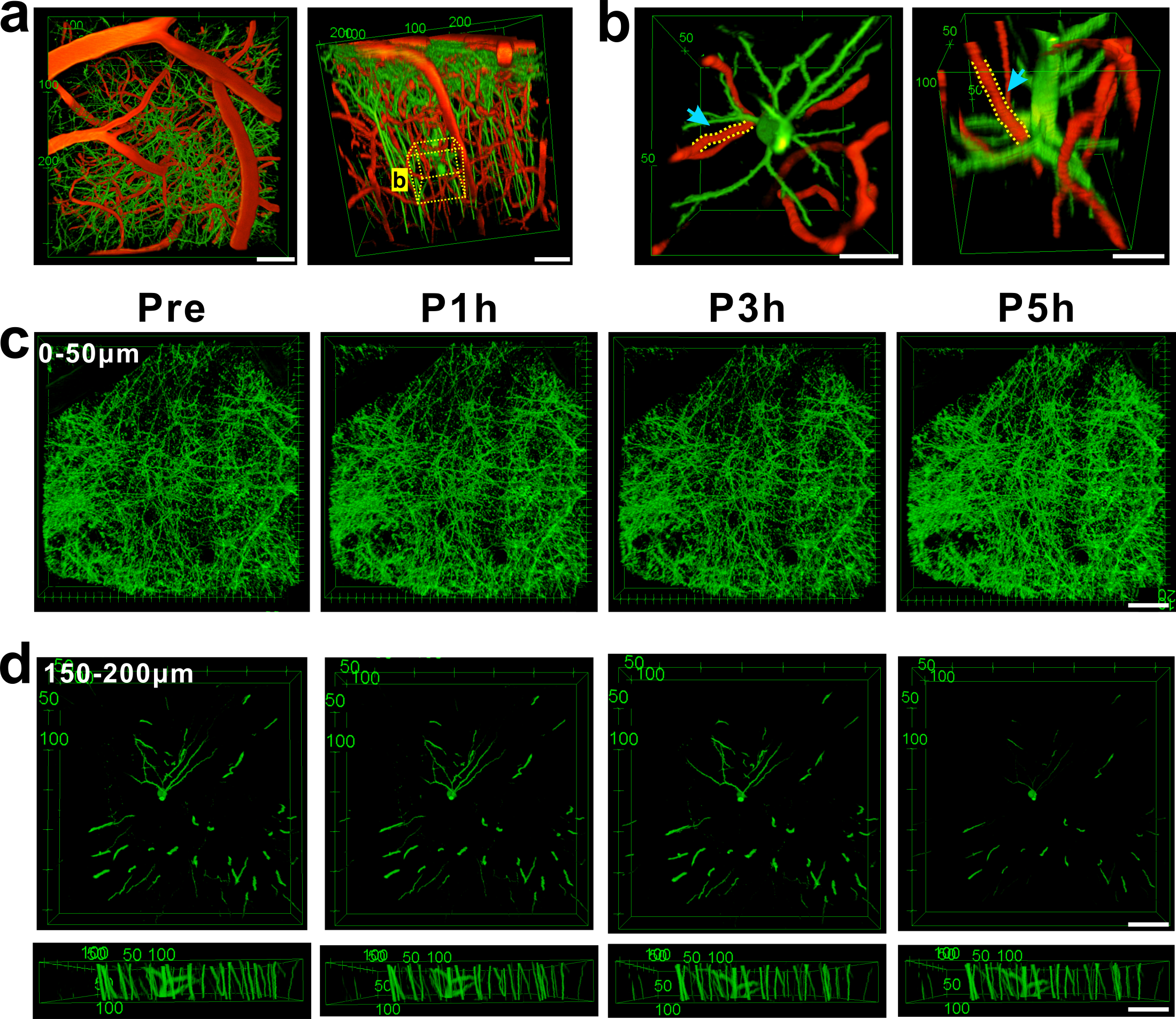
